# Following the track: accuracy and reproducibility of predation assessment on artificial caterpillars

**DOI:** 10.1101/2021.12.10.472105

**Authors:** Elena Valdés Correcher, Elina Mäntylä, Luc Barbaro, Thomas Damestoy, Katerina Sam, Bastien Castagneyrol

**Affiliations:** Univ. Bordeaux, INRAE, BIOGECO, F-33612 Cestas, France; Biology Centre of the Czech Academy of Sciences, Ceske Budejovice, Czech Republic; Faculty of Science, University of South Bohemia, Ceske Budejovice, Czech Republic; DYNAFOR, University of Toulouse, INRAE, Castanet-Tolosan, France; UniLaSalle, AGHYLE, 19 rue Pierre Waguet, 60026 Beauvais, France

## Abstract

Experimental studies of biotic interactions in real field conditions are essential to understand the structure and functioning of ecological networks. The use of artificial caterpillars to mimic actual prey availability is generally seen as a standard approach to compare the activity and diversity of predators along environmental gradients. Yet, even with standardized material and procedures, biases may still affect data from multiple observers with different expertise. We used pictures of artificial caterpillars with or without various predation marks, in an online survey that was targeted for the participants of the project, to evaluate the reliability of predation marks identification made by non-scientists and by scientists with and without previous experience in predation mark identification. Pictures of artificial caterpillars displayed typical marks left by birds, mammals and arthropods, as well as non-predation marks (‘false positive’). 357 respondents scanned 7140 pictures of these pictures. Self-declared scientists were more confident and accurate in their observations than non-scientists, but the differences in correct identifications among scientists and non-scientists were low. Self-declared scientists with previous experience were also more accurate than scientists without previous experience, while there were no differences in self-confidence among scientists with and without previous experience. Accuracy in predation mark identification did not differ among types of predators, but respondents were more keen to identify marks left by birds or mammals than arthropods. Our results have practical implications for the design of multi-observer projects relying on artificial caterpillars as a proxy to assess predation intensity, in particular in the context of citizen science.

## Introduction

By initiating top-down trophic cascades, predators can indirectly control the amount of plant biomass consumed by insect herbivores in both natural and agricultural landscapes (Vidal & Murphy, 2018; Abdala-Roberts et al., 2019). As a key biotic interaction, predation has been an ecological process scrutinized by ecologists for decades (Holmes et al., 1979; Fowler & Knight, 1991; Mäntylä et al., 2011). Yet, predation is a fleeting phenomenon that is difficult to track in real time, especially because it has delayed effects on prey demography and plant response. Direct observations provide unambiguous proof of predation, but they require spending long hours in the field (Harvey & Gittleman, 1992). The use of automated camera traps partially solved this issue, but although prices are dropping, this technology remains expensive and appropriate mostly only for large predators and prey (O’brien & Kinnaird, 2008; Muiruri et al., 2016; Akcali et al., 2019; Iannarilli et al., 2021). An alternative is to focus on the outcome of predation, rather than on predation itself. An approach in ecology consists of monitoring the biomass of primary producers in presence or absence of secondary consumers — *aka* herbivores’ natural enemies (Mooney et al., 2010). For instance, monitoring insect herbivory or plant growth while preventing vertebrate predators’ access to plants reveals the ecological importance of predators (Mäntylä et al., 2011). However, such exclusion experiments do not allow assessing the identity or functional diversity of the predators themselves, which is central to the understanding of predator-prey relationships involved in wider trophic cascades (Philpott et al., 2009; Maas et al., 2015). In addition, they do not quantify predation *per se* (Zverev et al., 2020).

Among the several other approaches used by ecologists to study predation, the deployment of artificial caterpillars as “sentinel” has bloomed in recent decades (e.g. Howe et al., 2009; Lövei & Ferrante, 2017; Rößler et al., 2018). In particular, many studies have used dummy caterpillars to comparatively assess predation rates across habitats or along natural or anthropic ecological gradients (Mäntylä et al., 2008; Barbaro et al., 2012; Tvardikova & Novotny, 2012; Sam et al., 2015; Valdés-Correcher et al., 2021). The method offers several advantages: it is easy to implement, replicate and standardize, it is relatively fast, and it has a very limited cost as compared to other methods. Several authors have designed experiments to match marks on artificial caterpillars with the identity of predators (Low et al., 2014; Sam et al., 2015; Khan & Joseph, 2021). However, whether a given mark can be attributed with high probability to an arthropod, a bird, a mammal, or a lizard, remains difficult and may stay uncertain in many instances. Such an uncertainty impedes a proper understanding of predation patterns and makes estimations of predation intensity experienced by local prey communities unreliable. Although the deployment of artificial caterpillar in the field can be done by citizens with no previous scientific expertise (Castagneyrol et al., 2020), standardizing the interpretation of marks should be of particular concern when predation marks are assessed by multiple observers across multiple sites along large geographical gradients (Roslin et al., 2017; Zvereva et al., 2020).

Still, the use of sentinel larvae offers an unprecedented opportunity to standardize the study of predation over large geographic areas and ecological contexts (Lövei & Ferrante, 2017). This is of particular interest in citizen science, or the volunteer contribution of non-professional scientists to the production of scientific knowledge (Dickinson et al., 2010; Valdés-Correcher et al., 2021). Considerable efforts have been made to understand, control and reduce sources of errors in observations made by citizen scientists (Ratnieks et al., 2016; Swanson et al., 2016; van der Wal et al., 2016) in order to improve the reliability of data (Kosmala et al., 2016; Balázs et al., 2021). For example, whether the use of sentinel larvae could be successful in citizen science programs is a matter of debate (Castagneyrol et al., 2020). Given the great scientific and pedagogical potential of sentinel larvae as a way to study predation and teach ecology (Curtis et al., 2013; Leuenberger et al., 2019), it is important to evaluate the reliability of the method when used by non-experts in the case that technical or financial constraints prevent shipping the material to a single expert. In this study, we developed a picture quiz to evaluate the accuracy and reproducibility of the identification of marks left on the surface of artificial caterpillars. We expected that self-declared scientists would be more confident with their identification, and more accurate than non-scientists, and also that self-declared scientists with previous experience in predation mark identification would be also more confident and accurate than scientists without previous experience. We further used the data to identify which type of marks are more likely to be misidentified by observers; we predicted that beak marks left by birds would be more often correctly identified than more discrete marks left by arthropod mandibles. By doing so, our study evaluates and discusses the reliability of predation rates assessments by experts and non-experts in citizen science programs using artificial caterpillars.

## Methods

### Tree bodyguards citizen science project

We recently initiated the *Tree bodyguards* citizen science project using artificial caterpillars to study trophic interactions in oaks across a large geographic gradient (Valdés-Correcher et al., 2021). The project involved both ecologists — who may or may not have worked with artificial caterpillars in the past — and school children and their teachers in 23 countries in Europe. We learnt from the first two years of the project that predation rates estimated by school children were significantly biased compared to scientists’ assessments (Valdés-Correcher et al., 2021). Interviews with teachers revealed that they had difficulties in teaching their pupils how to identify predation marks, as they themselves lacked significant field experience (Perron, 2021). We therefore developed a picture quiz as a training material to get teachers and pupils familiar with the various traces likely to be observed on artificial caterpillars.

### Designing picture quizzes

We used artificial caterpillars that have been exposed for 15 days on oak trees between May and June of 2018 and 2019 by the partners of the *Tree bodyguards* citizen science project. We photographed a subset of caterpillars presenting typical marks left by birds, arthropods, or mammals (henceforth, *predation* marks), and caterpillars with nail or leaf scratches, bud or branch imprints (henceforth, ‘*false positive*’), as well as caterpillars with smooth surfaces (**Figure 1**). Pictures were taken with a smartphone with a 48 Mpx camera in a standardized way in terms of light and position. We did not aim to have high resolution pictures but instead we used pictures comparable to those possibly made by project partners in the- field, that would be exchanged with scientists to confirm predation diagnostics.

**Figure 1:**
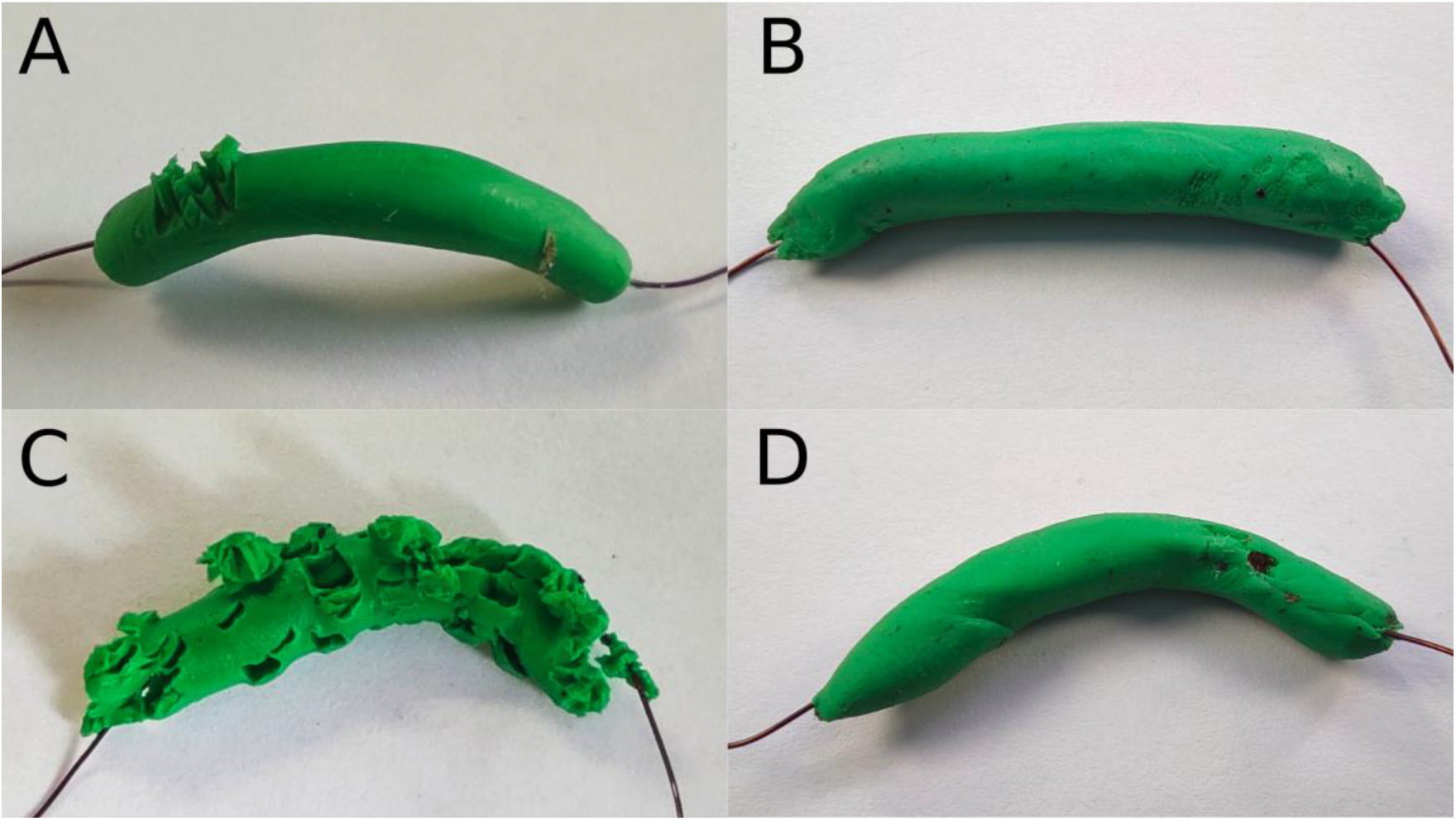
*A sample of pictures representing fake caterpillars attacked by birds (A), arthropods (B), mammals (C), or not attacked at all, but with marks left by branches (*false positive, *D).*

We assumed that we correctly identified predation marks. We double checked predation marks by several experts and also compared the pictures with published pictures or photographic keys such as the one in Low et al 2014. We acknowledge that this is questionable. However, we consider that having caterpillars at hand allowed us to make a more precise diagnostic than what respondents could do based on pictures.

We prepared three different picture quizzes with the same proportion of pictures of artificial caterpillars with bird (25%), arthropod (23%) or mammal (5%) predation marks, as well as pictures of caterpillars with smooth surface or marks considered as ‘false positive’ (47%). The proportions of each damage type was representative of what we observed in the field. The three quizzes presented different pictures in different orders.

We prepared four language versions of the three quizzes (English, French, German and Spanish). For a given quiz (1-3), the pictures were the same, and in the same order, in the different language versions. For each picture, we invited respondents to scrutinize the photographed caterpillar to identify potential predation marks. There were five possible choices: bird attack, arthropod attack, mammal attack, no attack or false attack, plus another “*I don’t know*” category (Unidentified marks) and the respondents had to select one of them.

We first distributed the quiz among the *Tree bodyguards* project partners in the monthly newsletter, and advertised it on social media (*Twitter*). In addition to the identification of marks on artificial caterpillars, we asked respondents to declare whether they were scientists, students, school children or teachers. We pooled the last three categories into a single one (non-scientist). We also asked scientists if they had previous experience in the identification of marks on artificial caterpillars. We aggregated responses across the different language versions of the three quizzes. Respondents were informed that the quiz was anonymous and that their answers will not allow them to be identified.

### Data analysis

#### Self-confidence

We defined *self-confidence* as the number of identified pictures. We considered a picture was identified as soon as respondents gave any answer other than “*I don’t know*”, regardless of whether the identification was correct or not. We tested whether self-confidence varied between scientists and non-scientists, and between scientists with and without previous experience using a generalized linear model (GLM) with quasibinomial error distribution (to account for overdispersion) and logit-link. The response variable was the number of identified pictures (excluding the “*I don’t know*” category) divided by the total number of pictures (percentage of assessed pictures). We considered Quiz ID (1, 2 or 3) and self-declared respondent type as explanatory factors.

We further asked whether the probability respondents declared that they could not identify predation marks varied among types of predation marks with a *χ*^2^ test of independence.

#### Accuracy

We then focused only on pictures that respondents identified (*i.e.* excluding the “*I don’t know*” category) and defined *accuracy* as the percentage of correct identifications for each picture. We tested whether consistency differed between scientists and non-scientists and between scientists with and without previous experience among types of predation marks using a generalized linear mixed effect model (GLMM) with binomial error distribution and logit-link. The response variable was binomial and consisted in the number of correct *vs* erroneous identifications. Explanatory factors were the self-declared category of respondent (*i.e.*, scientist *vs.* non-scientist) and the type of predation marks (birds, mammals, arthropods, none). We added respondent ID (*i.e.*, each participant had an ID that changed in case they repeated a quiz) and picture ID as partially crossed random factors to account for multiple responses made by each respondent and repeated assessment of the same picture.

Finally, we qualitatively explored misidentification at the level of each picture in order to get insights into which type of marks was more likely to be confused with another.

We conducted all analyses in the *R* environment for statistical computing (R Core Team, 2020) using libraries lme4 (Bates et al., 2018), car (Fox and Weisberg, 2019) and DHARMa (Hartig 2021).

## Results

We uploaded the results on 2021-07-16. The three quizzes totalled 357 answers (260 in French, 78 in English, 3 in Spanish, 16 in German). Of the 65 (*i.e.*, 18.2%) respondents that were self-declared scientists, 39 had no previous experience in the identification of predation marks. Of the respondents, 31 declared they participated in the *Tree bodyguards* citizen science project. This represented 49% of self-declared scientists who filled the questionnaire, and 27% of non-scientists. The latter result has to be considered with caution, as it cannot be excluded that children participating in the project were unaware of it.

### Self-confidence

Respondents examined 7140 pictures. Self-confidence was on average 94%. It was less variable and slightly greater in self-declared scientists than in non-scientists (*df* = 1, *F* = 4.57, *P* = 0.033, **Figure 2 A**) whereas it did not vary among scientists with and without previous experience in predation mark identification (*df* = 1, *F* = 0.52, *P* = 0.469, **Figure 2 B**). Self-confidence did not vary significantly among quizzes (*df* = 2, *F* = 2.65, *P* = 0.072).

**Figure 2:**
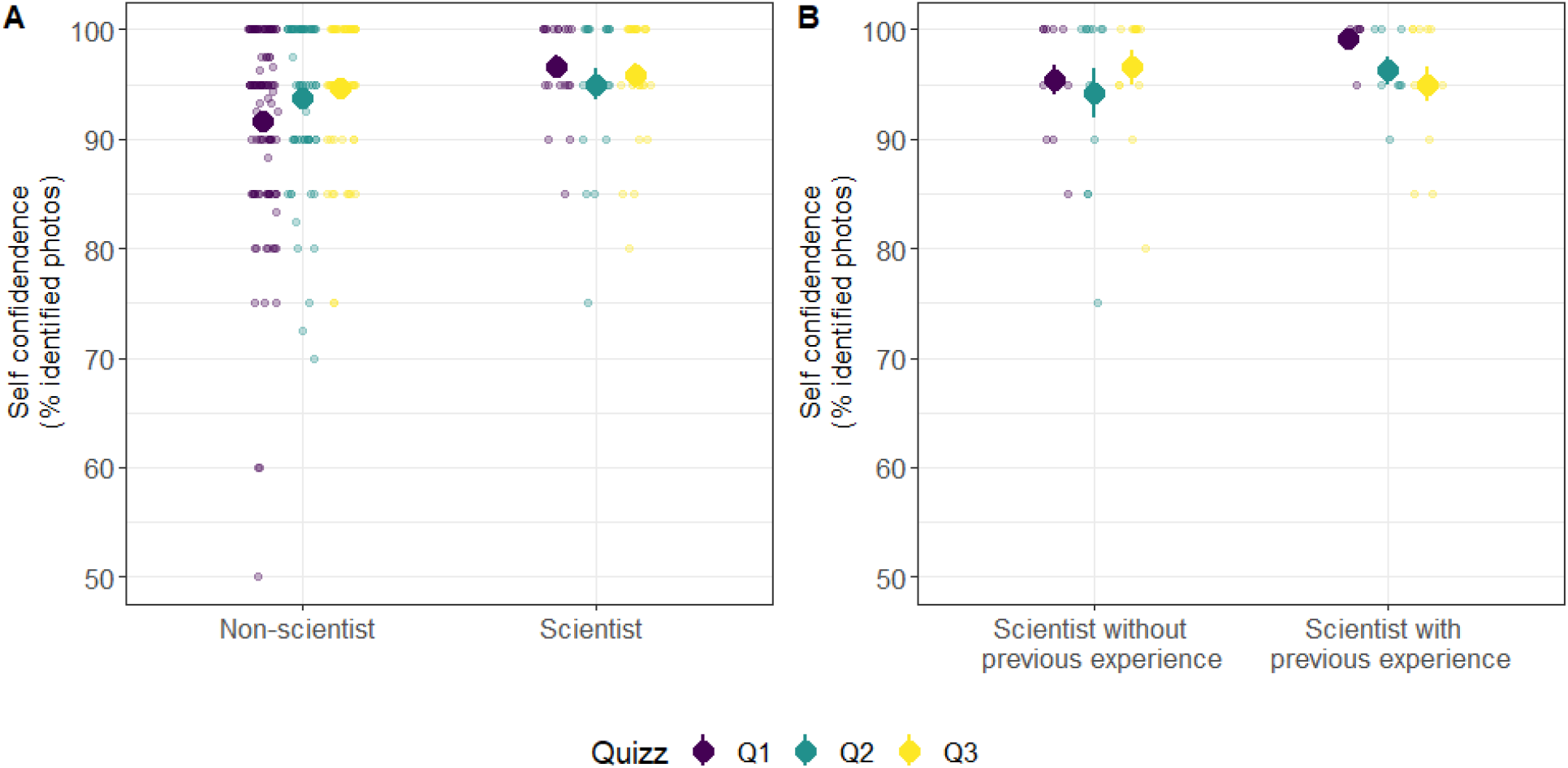
Self-confidence of self-declared scientists and non-scientists (A) and of scientists without and with previous experience in predation mark identification (B). Self-confidence corresponds to the percentage of pictures that respondents identified, regardless of whether the identification was correct or not. Small transparent dots represent raw data. Large dots and corresponding error bars represent raw means and standard errors.

Of the 456 pictures that respondents declared as unidentifiable, the majority (57%) represented caterpillars with no predation marks or caterpillars with marks left by oak leaves, buds or branches; others represented caterpillars with arthropod marks (28%) or bird (12%) or mammal attacks (2%) **(Figure 3)**. *χ*^2^ test of independence indicated that the proportion of pictures that respondents could not identify varied significantly among the types of marks on caterpillars (*χ*^2^ = 0.12, *P*-value = 0.010). For sake of illustration, **Figure 4** shows the top-four pictures respondents could not identify.

**Figure 3:**
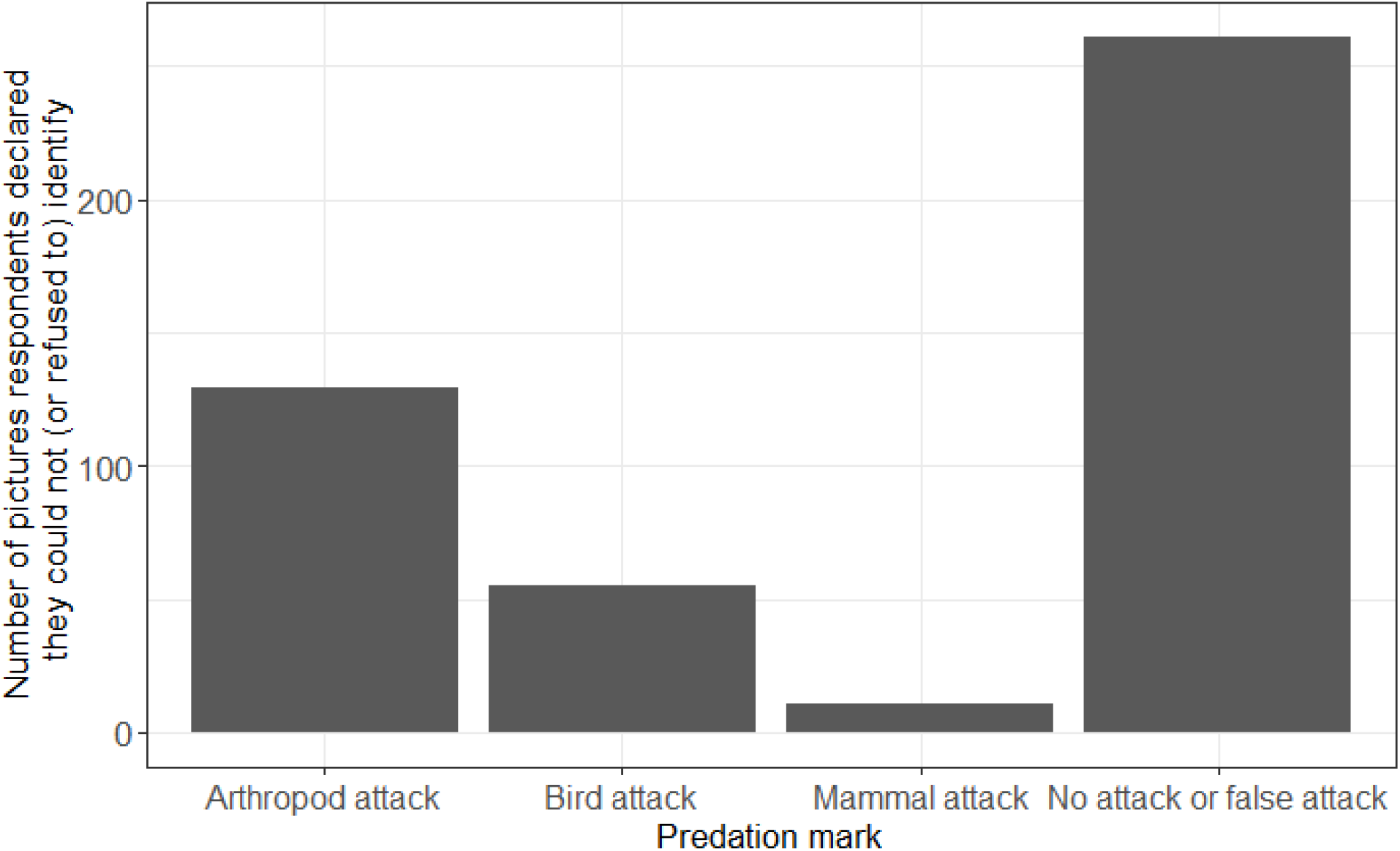
Number of pictures of each type of predator attack that respondents declared they could not (or refused to) identify.

**Figure 4:**
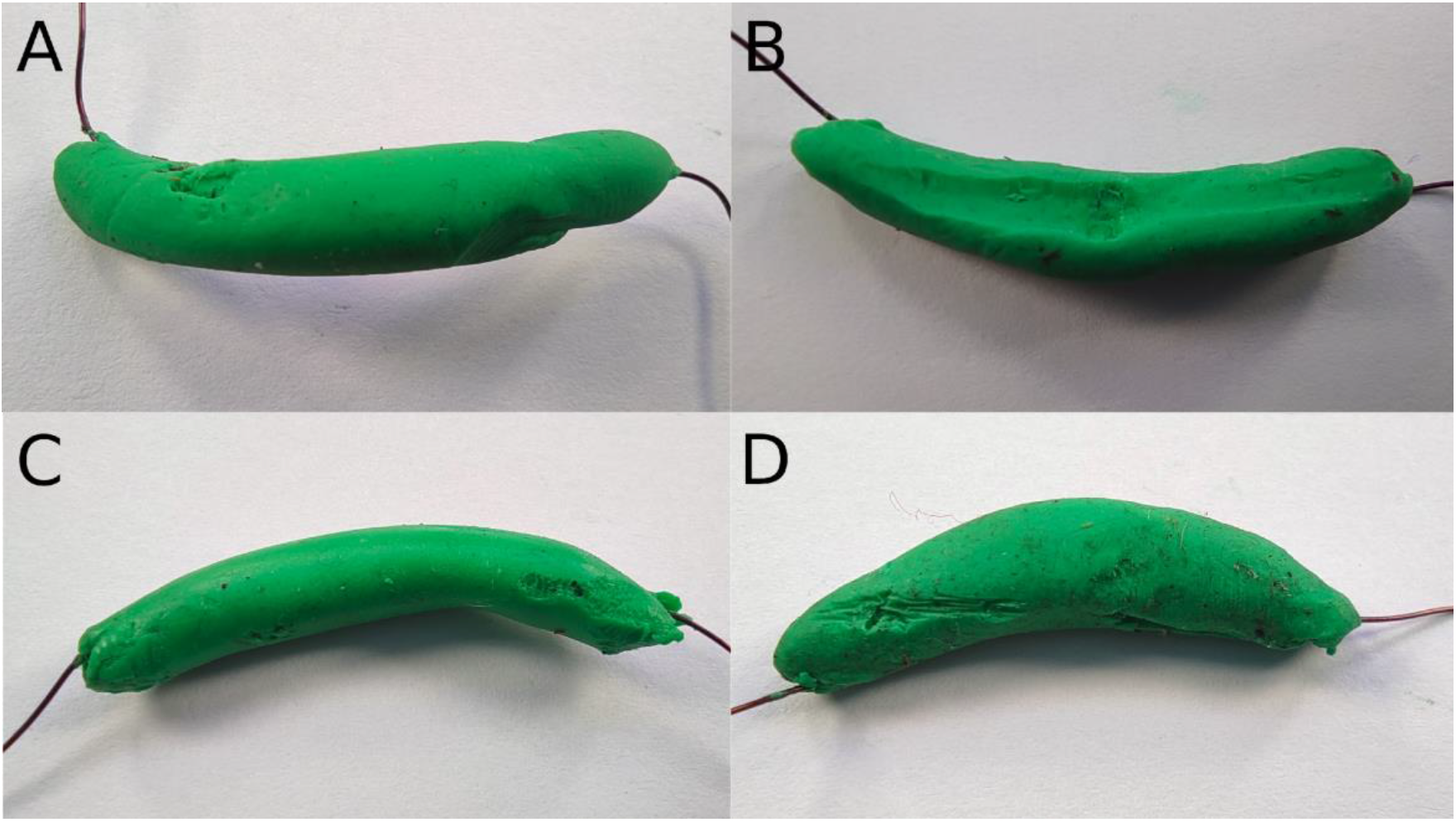
The four pictures which were declared as unidentifiable most frequently by respondents. A, B and D represent caterpillars with no predation mark (i.e. imprints of branches), and C a caterpillar with evidence of arthropod attacks.

### Accuracy

Accuracy of correct identifications for each picture was on average 70%. It was significantly higher in scientists (76 %) than in non-scientists (68 %) (df = 1, *χ*^2^ = 21.1, *P* < 0.001, **Figure 5 A**), and in scientists with previous experience (79 %) than in scientists without previous experience (74 %) in predation mark identification (df = 1, *χ*^2^ = 3.98, *P* = 0.046, **Figure 5 B**). Accuracy did not differ significantly among types of predation marks (df = 2, *χ*^2^ = 1.44, *P* = 0.697). It is however worth noticing that mismatches between identifications made by experts *vs* respondents were more frequent when dummy caterpillars had no predation marks, or arthropod predation marks **(Figure 6)**: respondents failed to recognize arthropod marks and considered them as ‘false positive,’ or conversely attributed non-predation marks to arthropod marks.

**Figure 5:**
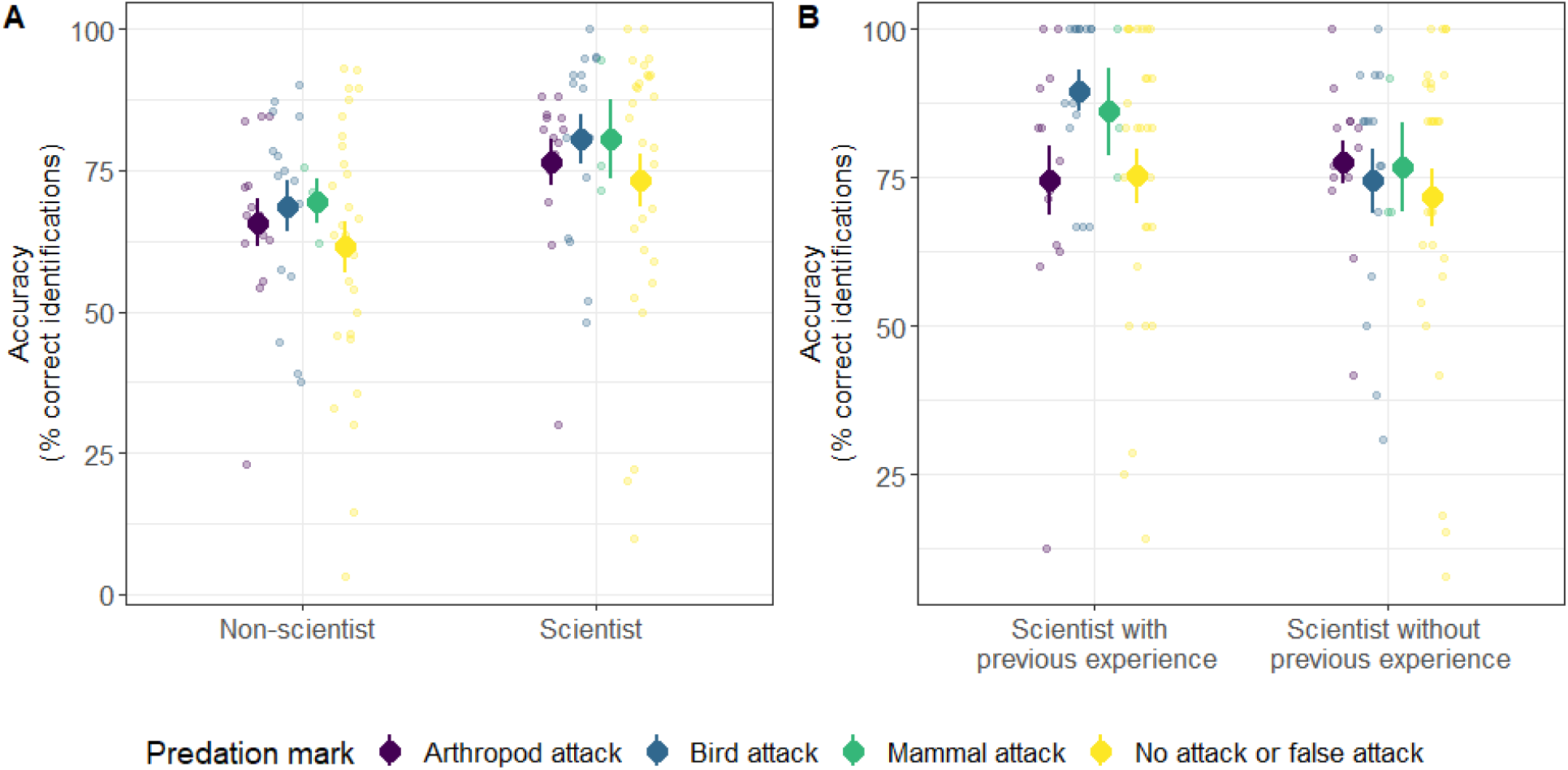
Accuracy of predation mark identification by self-declared scientists and non-scientists (A) and by scientists with or without previous experience (B). Accuracy corresponds to the percentage of pictures that respondents identified consistently with what experts did. Small transparent dots represent raw data. Large dots and corresponding error bars represent raw means and standard errors.

**Figure 6:**
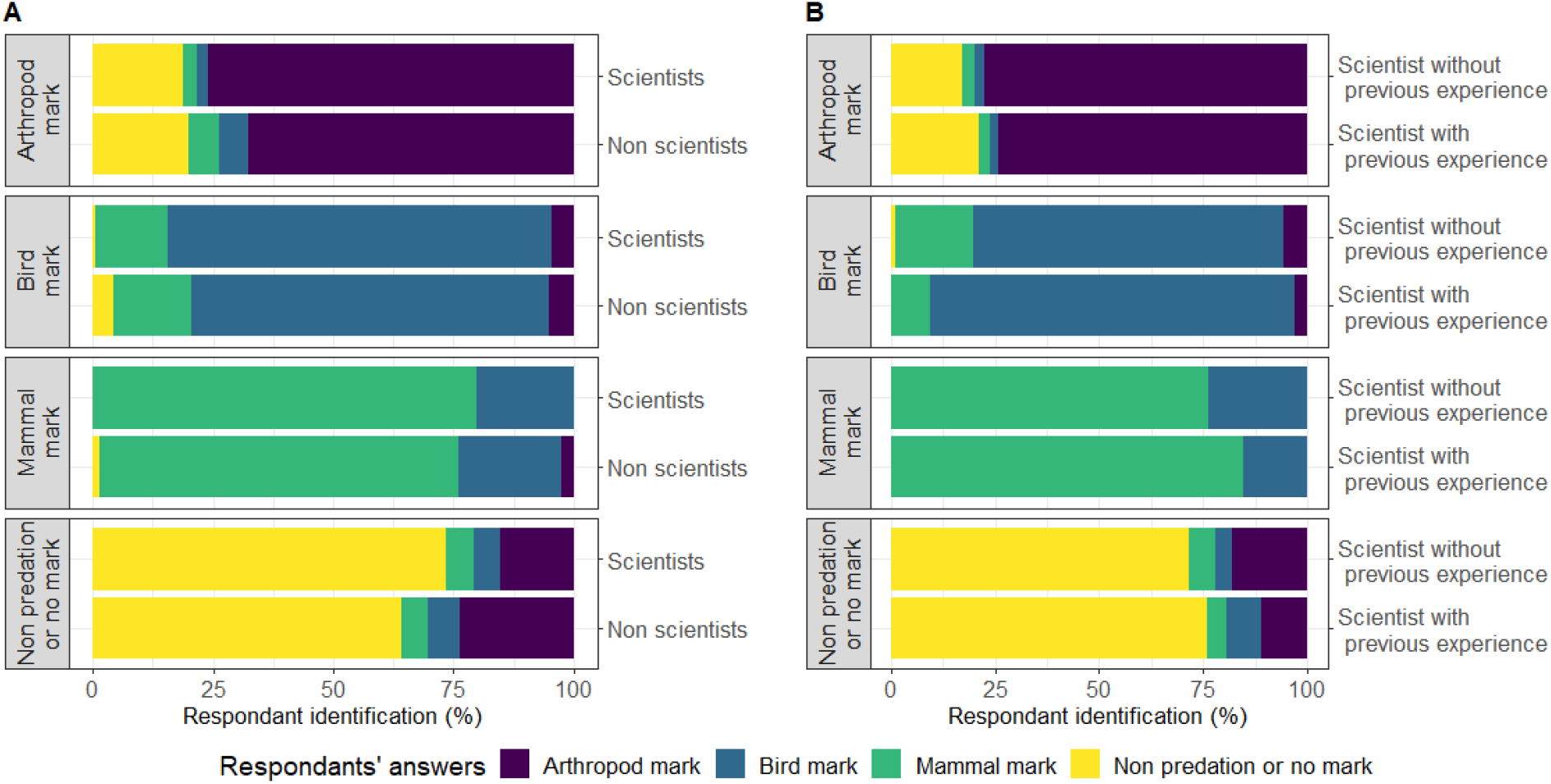
Comparison of expert’s and respondent’s identification of predation marks, according to respondents’ expertise. Expert identification is on the y-axis, in different panels. Colours refer to respondents’ identification. A: For each type of mark identified by experts, identification by self-declared scientists is compared with identification by non-scientists. B: For each type of mark identified by experts, identification by self-declared scientists with previous experience in predation rate assessment is compared with identification by self-declared scientists with no previous experience.

## Discussion

The use of artificial caterpillars is generally regarded as an appropriate standard approach to compare the activity of predators along ecological gradients (Howe et al., 2009; Lövei & Ferrante, 2017; Rößler et al., 2018). Yet, covering large ecological gradients generally requires extensive collaborations between multiple observers gathered in scientific networks, who might have different degrees of expertise for a specific skill or another (Roslin et al., 2017; Zvereva et al., 2020; Valdés-Correcher et al., 2021). This may cause uncontrolled biases in the estimation of predation rates, that have surprisingly seldom been investigated to date (Rößler et al., 2018). We found that the self-confidence and accuracy in the identifications of predation marks on artificial caterpillars, although it was high in every case, varied among self-declared scientists and non-scientists, and among scientists with and without previous experience in predation mark identification.

### Self-confidence

Self-declared scientists were more confident with their identifications of predation marks on artificial caterpillars than non-scientists were, whereas there were no differences among scientists with and without previous experience in predation mark identification. This result is somewhat surprising as our questionnaire invited the answer “*I don’t know*” and scientists should be more aware of the importance of a conservative scoring. In addition, self-confidence in one’s own skills often decreases with expertise (*aka* the “Dunning-Kruger effect,” after Kruger & Dunning, 1999). We therefore expected that self-declared scientists would have been more reluctant than less experienced respondents to classify predation marks in case their identification was ambiguous. A likely explanation is that some self-declared scientists had previous field experience with the use of artificial caterpillars. Besides, both self-declared experienced and non-experienced scientists may have taken the quiz more seriously and spent more time on the choice of the answer as they are used to perform scientific experiments (Johnson et al., 2016), which may also explain why they were also more accurate.

Interestingly, the comparison among types of marks on artificial caterpillars showed that respondents were more confident in identifying marks left by birds or mammals, and that these marks were consistently also more often correctly identified. Birds and mammals generally leave deep marks in artificial caterpillars, where the ‘V’-shaped mark left by bird bills or parallel lines left by rodent teeth are generally clearly visible (Low et al., 2014). On the contrary, arthropods can impress a greater diversity of shallower marks that are much more difficult to identify with certainty (Khan & Joseph, 2021). Arthropod marks can be as varied as scratches, pricks, granulated or disturbed surfaces (Khan & Joseph, 2021) that are often difficult to decipher from impressions left by buds, branches or leaves. Our study therefore confirms the expectation that respondents would identify bird and mammal attacks more accurately. Together with concerns that have been emitted regarding the biological significance of arthropod marks on artificial caterpillars as well as their potential variation under various climatic conditions (Rößler et al., 2018; Muchula et al., 2019; Khan & Joseph, 2021), these results call for caution when interpreting large-scale variability in predation rates by arthropods (Roslin et al., 2017; Zvereva & Kozlov, 2021).

### Accuracy

Self-declared scientists were more accurate in their identifications than non-scientists (Castagneyrol et al., 2020), and scientists with previous experience were also more accurate than scientists without previous experience (Johnson et al., 2016). Scientific expertise is not restricted to professional scientists, and research on participants in citizen science projects demonstrated that taxonomic expertise can be comparable between professional scientists and part of the general public (Austen et al., 2016, 2018). However, artificial caterpillars are not something the general public can be familiar with, and self-learnt identification skills are unlikely in this case. The better performance of self-declared scientists in general, as well as self-declared scientists with previous experience, therefore did not come as a surprise. Still, the percentage of agreement in their identifications as compared to that made by a single expert was 76% for self-declared scientists and 79 % for self-declared scientists with previous experience. Should they have had the artificial caterpillar at hand, it is likely the percentage of agreement would have been higher. Even though pictures were focused on damages on plasticine surface, we acknowledge that the quality of pictures as well as the resolution of screen devices on which they were displayed may have prevented accurate identification of marks on the surface of artificial caterpillars. We therefore recommend that predation marks are assessed with the caterpillar in hand as it allows seeing the scale and all the dimensions of the caterpillar, and also using a magnifier lens for small marks.

Non-scientists were slightly less accurate in their identifications than scientists (only 8% difference of accuracy) and it was also the case for scientists without previous experience in relation with scientists with previous experience. However, the accuracy of non-experts was in the range of what was found in previous studies. For instance, Low et al. (2014) reported that scientists had 68% of accuracy in their identification of different types of predation marks (birds, mammals and arthropod marks). Focusing on species identification, Khan and Joseph (2021) found that non-expert volunteers (including graduate students, laboratory technicians, post-docs and faculty members) had 85% of accuracy identification of common turfgrass arthropods. It is important to highlight that the differences in accuracy among scientists and non-scientists and among scientists with and without previous experience that we found, although significant, were pretty low, suggesting that both scientists and non-scientist can actually provide data accurate enough to support ecological research. We suggest that the use of a photographic reference collection to refer to can help identify the most typical marks on artificial caterpillars.

## Conclusion and practical implications

Several authors proposed guidelines for estimating predation rates on artificial caterpillars across ecological gradients, and recommended procedures to standardize protocols (Howe et al., 2009; Low et al., 2014; Lövei & Ferrante, 2017; Rößler et al., 2018). In this study, we went a step further by demonstrating that although we confirm the high interest and overall reliability of the method, standardization attempts are not completely satisfying if predation rate is to be assessed by multiple observers (Castagneyrol et al., 2020). The reason was twofold: (i) accuracy in predation mark identification varied substantially among scientists and non-scientists, and among scientist with and without previous experience (although less than expected), and (ii) subtle impressions by arthropods remain difficult to identify, both by scientists and non-scientists, and can easily be confounded with false mark attacks. Knowing the limitation of a method is the first step towards its improvement. Whenever possible, we recommend that predation rate is assessed on raw material, with a magnifier lens, by a single skilled observer or a few trained observers confronting their observations (Low et al., 2014). If artificial caterpillars have to be shipped across long distances, it is crucial that they are secured in individual vials preventing any damage during transportation, and stored at temperatures that are not too hot to avoid melting (Muchula et al., 2019). If technical or financial constraints prevent shipping the material to a single expert, we recommend that every observer is guided and tested with a photographic reference collection that includes pictures of typical predation marks, as well as pictures of “false positives”. Of course, a training session including a set of artificial caterpillars with different predation marks is needed to increase their self-confidence and accuracy. Besides, we also recommend including several checks of data quality and appropriate mitigation procedures in order to avoid errors in the identification of predation marks.

## Acknowledgements

This study was permitted by the financial support of the BNP Paribas Foundation through its Climate & Biodiversity Initiative for the *Tree bodyguards* citizen science project. EM was supported by the Grant Agency of the Czech Republic (19-28126X). KS was supported by ERC Starting Grant BABE 805189. The authors warmly thank the scientists, teachers and students who took part in this survey.

## Data accessibility

Data used in this study is available on the INRAE data portal: Valdés-Correcher, Elena; Mäntylä, Elina; Barbaro, Luc; Damestoy, Thomas; Sam, Katerina; Castagneyrol, Bastien, 2021, “Original data -- Following the track: accuracy and reproducibility of predation assessment on artificial caterpillars”, https://doi.org/10.15454/V4FGJV, Portail Data INRAE.

